## supplementary material for "Activator-blocker model of transcriptional regulation by pioneer-like factors"

***contributed equally**

[2]Present address: Epigenetics and Neurobiology Unit, European Molecular Biology Laboratory, EMBL Rome, Adriano Buzzati-Traverso Campus, Via Ramarini 32, 00015 Monterotondo (RM), Italy

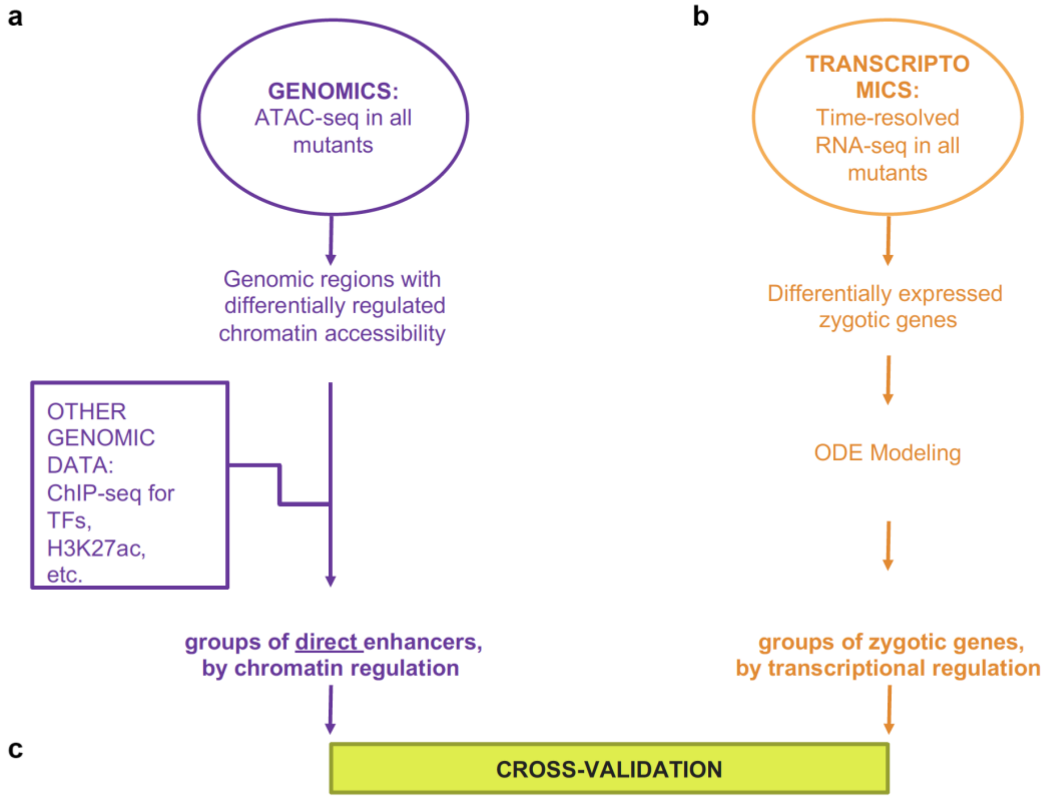

**Figure S1. Study design and outline of the data analysis (related to all figures).**
**a.** Genomic analysis. We performed Omni ATAC-seq in eight genotypes (WT, MZ*spg*, MZ*sox19b*, MZ*nanog*, MZ*ps*, MZ*sn*, MZ*pn* and MZ*triple*) and took 102 945 regions accessible in the wild-type (ARs) for further analysis. To select and group direct enhancers, we combined chromatin accessibility changes on ARs in each mutant with additional genomic information, which was the direct binding of TFs (ChIP-seq for Pou5f3, SoxB1, and Nanog) and changes in enhancer activation (ChIP-seq for H3K27ac active enhancer mark in the WT and selected mutants). This genomic analysis resulted in a list of direct enhancers bound by Pou5f3, SoxB1, or Nanog, classified by differential regulation. Data related to genome analysis are in the Table S1. **b.** Transcriptomic analysis. We performed time-resolved RNA-seq profiling at 8 developmental points, every 30 min starting from pre-ZGA (2.5 hpf) till midgastrulation (6 hpf), in 8 genotypes (WT, MZ*spg*, MZ*sox19b*, MZ*nanog*, MZ*ps*, MZ*sn*, MZ*pn,* and MZ*triple*), in at least two biological replicates per genotype. First, we derived differentially expressed gene groups (“DOWN”, “SAME”, “UP”) by pairwise comparisons of transcriptional profiles in each mutant to the wild-type. Second, to decipher combinatorial regulatory inputs of Pou5f3, SoxB1, and Nanog factors to transcription, we applied dynamic mathematical modeling to classify 1799 transcripts, downregulated in MZ*triple*, by type of regulation. This transcriptomic analysis resulted in a list of differentially regulated transcriptional targets of Pou5f3, SoxB1, or Nanog. Data related to transcriptome analysis are in the Table S2. **C.** Cross-validation. We put together the results of independent genome and transcriptome analysis parts to cross-validate them. We linked all zygotic genes to ARs within +/- 50 kb from the gene promoter, and computed correlations between enhancer and transcript groups (Table S3).

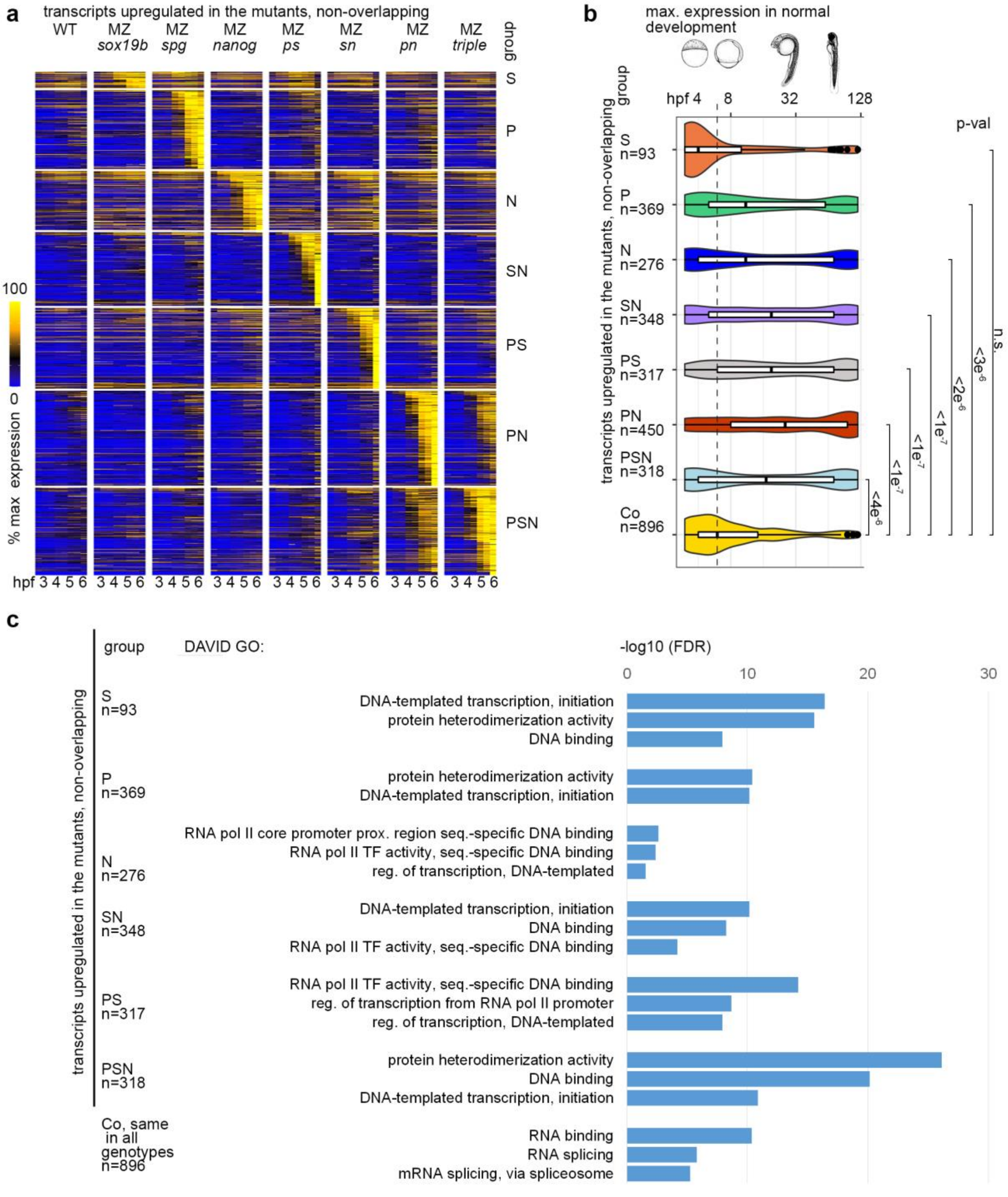

**Figure S2. DNA-binding proteins are prematurely expressed in most PSN mutants (related to the main Fig. 2).**

**a**. Heatmap of expression profiles of zygotic transcripts, upregulated in at least one mutant relatively to the WT, in all igenotypes. Transcripts were sorted to 7 non-overlapping groups by maximal expression (Table S2). Groups were named by absent TF(s) in the mutant (vertical labels). **b**. Premature transcription of zygotic genes occurs in all mutants except MZ*sox19b*. The max. expression in the time window from 3 hpf to 5 days (120 hpf) of normal development was calculated for each gene. Median expression time for each group was compared to the control group (Co): 896 zygotic transcripts expressed in the WT and not significantly changed in any mutants. 1-way ANOVA, p-val<$2e^{-16}$, p-values in Tukey-Kramer test are shown. **c**. DNA-binding proteins are prematurely expressed in the mutants for P, S, and N. DAVID, "Molecular function" GO analysis. Note the enrichment for DNA-binding proteins and regulatory function in transcription in upregulated groups, and enrichment for housekeeping genes in the unchanged control group. Source data are provided as a Source Data file 1. Zebrafish embryo drawings were used with permission of John Wiley & Sons - Books, from “Stages of Embryonic Development of the Zebrafish”, Kimmel et al., Developmental Dynamics 203:253-310 (1995); permission conveyed through Copyright Clearance Center, Inc.

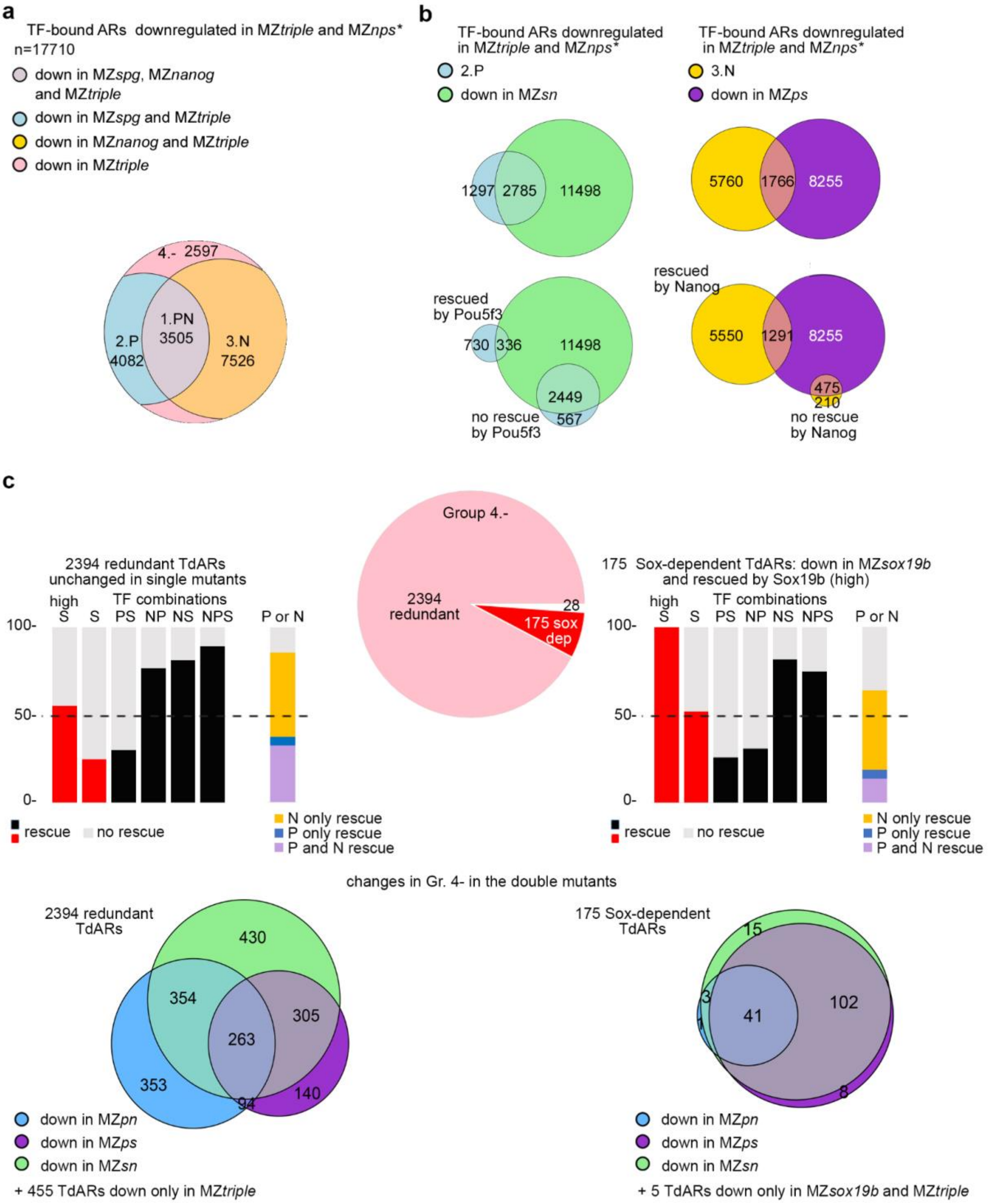

**Figure S3. Pou5f3, Sox19b and Nanog have different pioneer-like factor properties (related to the main Fig. 3).**

**a.** TdARs (TF-bound ARs, downregulated in MZ*triple*) which were also downregulated in the triple mutant MZ*nps* (Miao et al., 2022)[5] were divided to 4 groups by non-redundant requirements for Pou5f3 and/or Nanog for chromatin accessibility (as in Fig. 3a, main text). **b.** top: Venn diagrams show the overlaps between the 2.P and 3.N groups of TdARs, downregulated in the respective single mutants, with TdARs, downregulated in the double mutants by the other two factors. Bottom: The same as top, but 2.P and 3.N groups were split to two groups by rescue by Pou5f3 or Nanog, respectively, in MZ*nps*[5]. Left: Pioneer-like activity of Pou5f3 requires assistance of SoxB1 or Nanog in the majority of cases. 59% of 2.P group were downregulated in Sox19b/Nanog double mutant MZ*sn* (top); out of them 31% of 1066 regions which could be rescued by Pou5f3 in MZ*nps* and 82% of 3016 regions which could not be rescued by Pou5f3. Right: Pioneer-like activity of Nanog does not require assistance of SoxB1 or Pou5f3 in the majority of cases. Only 23% of 3.N group were also downregulated in Pou5f3/Sox19b double mutant MZ*ps* (top); out of them 19% of 6841 regions which could be rescued by Nanog in MZ*nps* and 69% of 685 regions which could not be rescued by Nanog. **c.** Analysis of TFs redundantly required for chromatin accessibility in Group 4.- TdARs**.** Pie chart (pink), left part of the panel: 2394 regions were redundant (not downregulated in any single mutant). Pie chart (red) and the right part of the panel: 175 regions were Sox19b-dependent: downregulated in MZ*sox19b* and rescued by high Sox19b concentration. 28 TdARs (downregulated. In MZ*sox19b* and not rescued by Sox19b) were omitted. Middle row: rescue experiment[5], Pou5f3 and Nanog rescue shown for high concentration. Bottom row: Venn diagrams of downregulated TdARs in the double mutants. Source data are provided as a Source Data file 1.

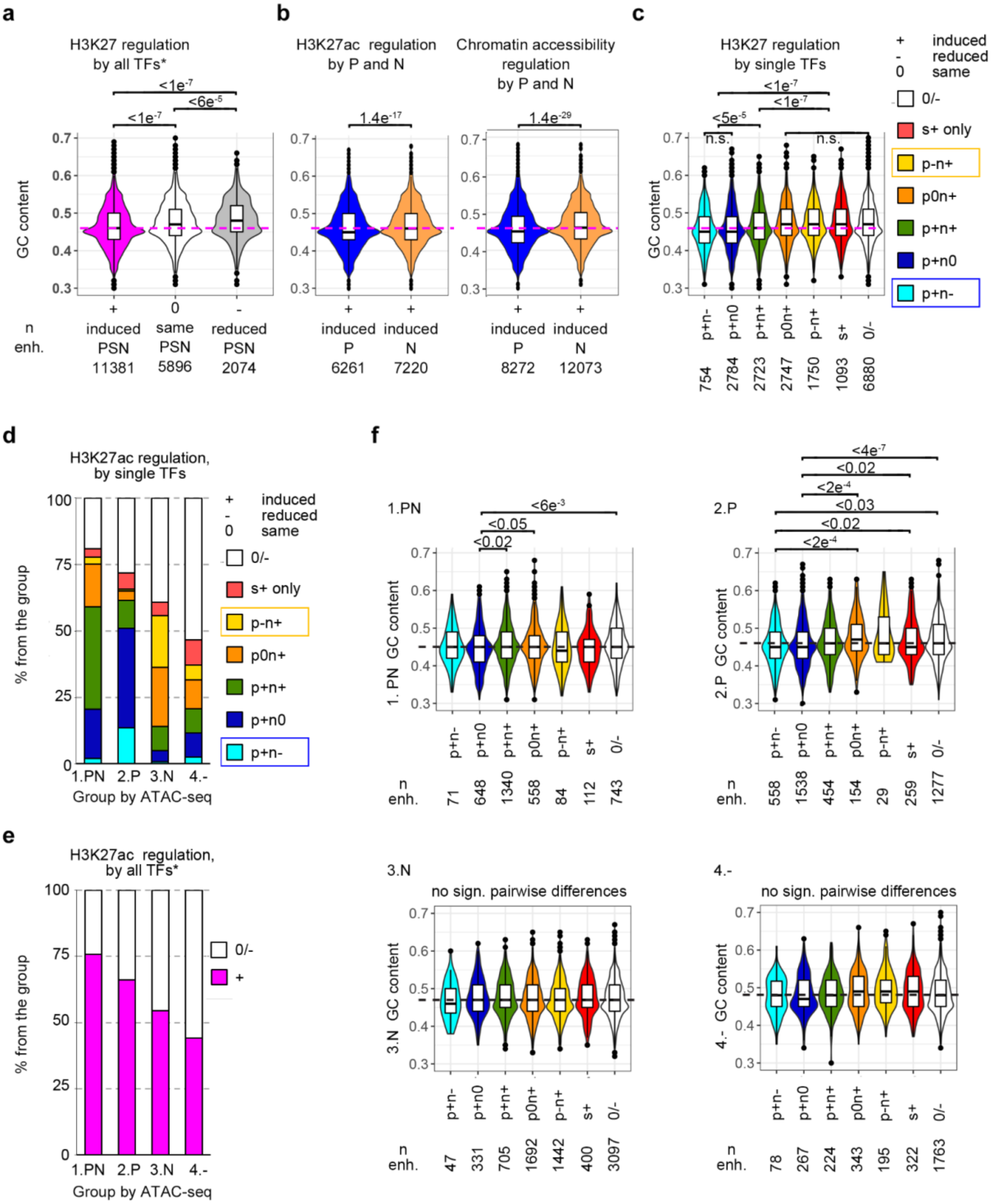

**Figure S4. Enhancer classification by TF effects on H3K27ac (related to the main Fig.4).**
**a-c, f**. GC content of indicated enhancer groups. TdARs H3K27-acetylated in any genotype were scored as enhancers. Dashed magenta line shows the median GC level of enhancers where H3K27ac was induced by combination of the factors (PSN, * H3K27ac data from [5]). P-values for statistically significant pairwise differences are shown in Tukey-Kramer test (a, c, f) or in Student 2-tailed t-test (b). **a.** 1-way ANOVA p-val < $2e^{-16}$ **b.** Pou5f3 induces H3K27ac (left) and creates open chromatin regions (right) in the regions with significantly lower overall GC content than Nanog. The P and N groups were partially overlapping but still significantly different in GC content **c.** Enhancer classification by the effects of the single TF combinations on H3K27ac. H3K27ac can be induced (+), unchanged (0), or reduced (-) by each TF. Antagonistic enhancers p+n- and p-n+ are boxed in the legend. 1-way ANOVA p-val < $2e^{-16}$ **d.** Percentages of enhancer types by regulation by single TFs shown in (c) within each of the groups 1.PN,

2.P, 3.N and 4.-. **e**. effects of PSN combined activity on H3K27ac within the groups 1.PN, 2.P, 3.N and 4.-. Note that enhancer activation by PSN does not exceed the sum of activation by single factors (compare to d). **f.** GC content of all the enhancer groups shown in (d). Note that subgroup 1.PN(p+n0) has the lowest GC content within 1.PN group; subgroups 2.P(p+n0) and 2.P(p+n-) had the lowest GC content within 2.P group. Dashed line – median GC content of 1.PN, 2.P, 3.N and 4.- groups. P-values in 1-way ANOVA were p= 0.000901 for 1.PN, p= 2.04e-11 for 2.P, p=0.0421 for 3.N, p=0.0127 for 4.- groups. Source data are provided as a Source Data file 1. The centers of the box plots correspond to the median values, the lower and upper bounds of the box correspond to the 25th and 75th percentiles, the upper whisker extends from the upper bound to the largest value no further than 1.5 * IQR ( inter-quartile range), the lower whisker extends from the lower bound to the smallest value at most 1.5 * IQR. Outlying points beyond the end of the whiskers are plotted individually.

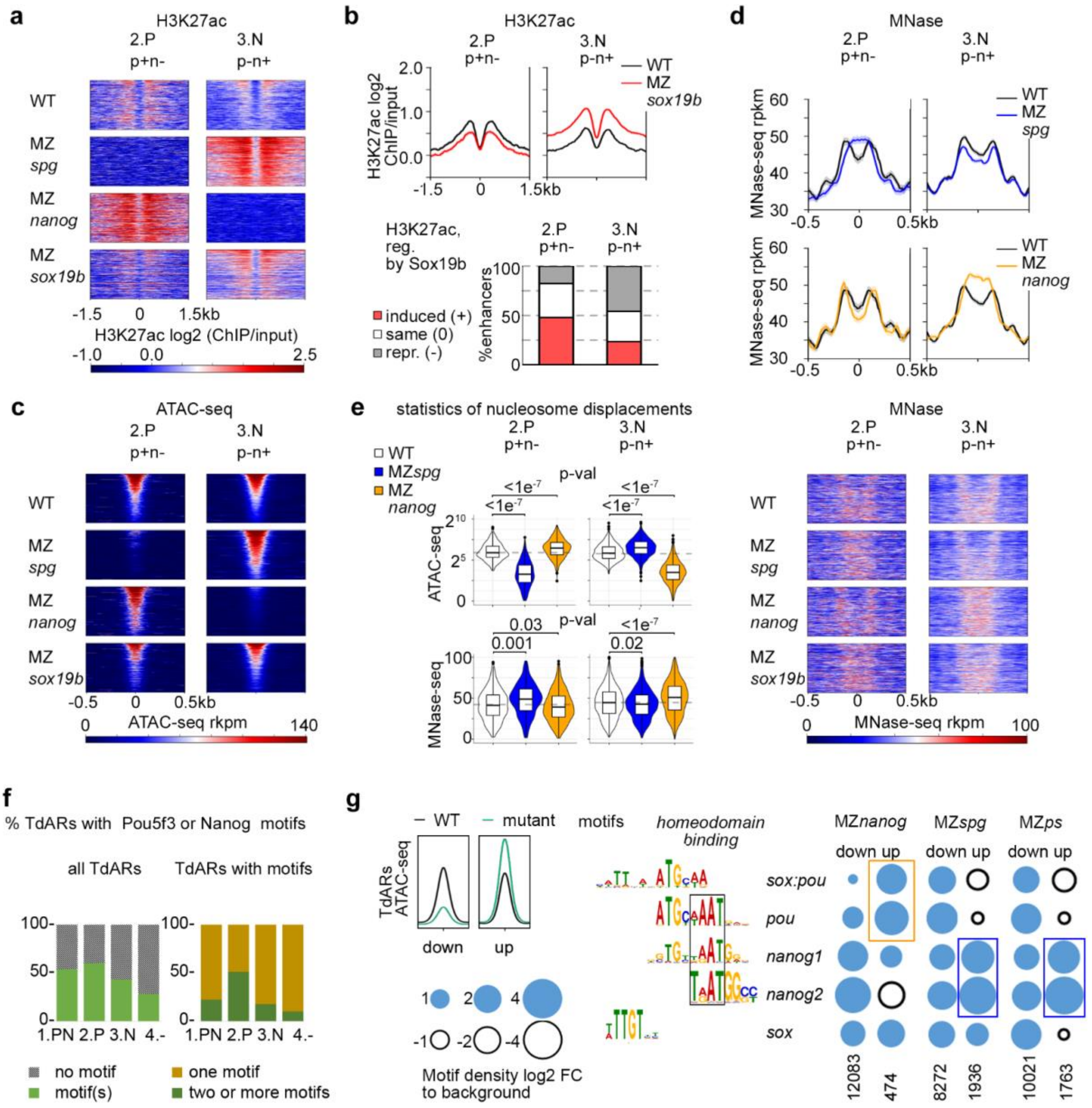

**Figure S5. Pou5f3 and Nanog compete as activator (+) and blocker (-) on antagonistic enhancers (related to the main Fig.4).**

**a.** Heatmap for H3K27ac in the indicated genotypes, on the 558 2.P p+n- and 1442 3.N p-n+ antagonistic enhancers. b**.** Top: summary profiles for H3K27ac in the WT and MZ*sox19b* on the 2.P p+n- and 3.N p-n+ enhancers (heatmaps in a). Bottom: Sox19b cooperates with Pou5f3 in more cases than with Nanog: **c**. Heatmaps for ATAC-seq in the indicated genotypes, on the 558 2.P p+n- and 1442 3.N p-n+ antagonistic enhancers. **d.** Summary profiles and heatmaps for MNase-seq in the indicated genotypes, for the 2.P p+n- and the 3.N p-n+ enhancers. In (a,c,d) enhancers were sorted by descending ATAC-seq signal in the WT. **e**. The opposite effects of Pou5f3 and Nanog on chromatin accessibility and nucleosome occupancy on the 2.P p+n- and 3.N p-n+ antagonistic enhancers are statistically significant. Violin plots show the distribution of ATAC-seq signal (top) and MNase-seq signal (bottom) in the WT, MZ*spg* and MZ*nanog*. P-values in Tukey-Kramer test are shown above the graphs. 1-way ANOVA, ATAC-seq p-val $<2e^{-16}$; 1-way ANOVA, MNase-seq p-val $=7.2e^{-9}$ for 2.P p+n- enhancers, p-val$<2e^{-16}$ for 3.N p-n+ enhancers. **f.** The numbers of merged motifs for Pou5f3 or Nanog (*sox:pou, pou, nanog1, nanog2)* in 110bp TdARs. **g.** TdARs downregulated (down) or upregulated (up) in each mutant compared to wild-type were selected to score enrichment of sequence -specific motifs (scheme at the left). Motif frequencies in down- and upregulated regions for the indicated genotypes; all TdARs - TF-bound accessible regions downregulated in MZ*triple*. Number of regions at the bottom. Note the enrichment for Pou5f3 cognate motifs in the regions upregulated in MZ*nanog* (yellow box) and enrichment for Nanog cognate motifs in the regions upregulated in MZ*spg* and MZ*ps* (blue boxes). Source data are provided as a Source Data file 1.

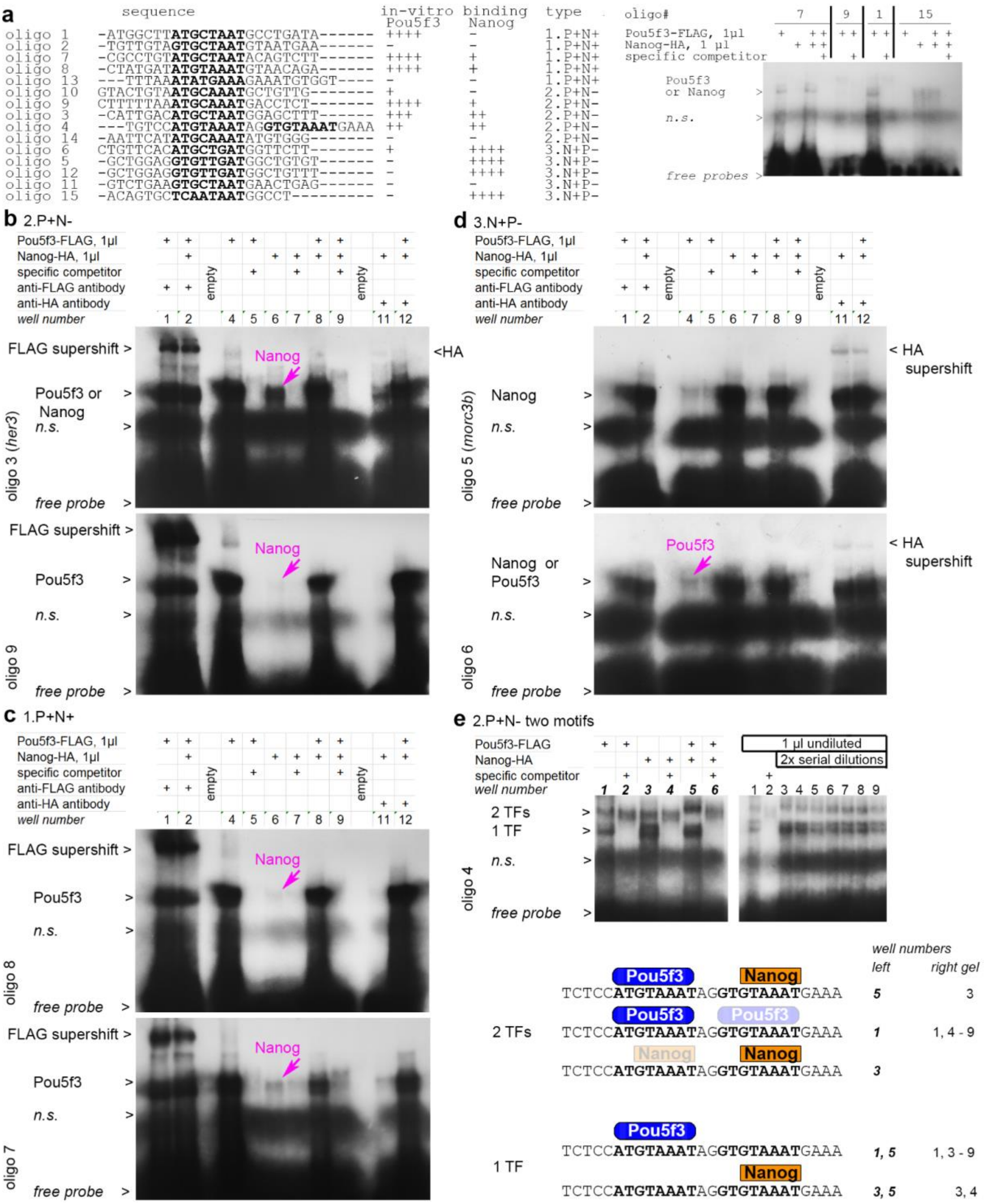

**Figure S6. Pou5f3 and Nanog bind to the common motifs with different affinity (related to the main Fig. 5).**

**a.** Five oligos were designed for each of the three types of enhancers, differing by TF effects on chromatin accessibility: 1.P+N+, 2.P+N-, 3.N+P-. Left: names, CLUSTAL-aligned sequences of the EMSA oligos (octamer sequences matching to Pou5f3 or Nanog motifs are in bold), and the summary of the results. Right: Autoradiogram with 3 hr exposure shows strong binding (++++) of Pou5f3 to oligo 7 and 1, and of Nanog to oligo 15. Moderate binding (++) of oligo 9 is hardly detectable with this exposure time. **b-d**. EMSA for the indicated Nanog- and Pou5f3- binding oligos with single motifs, long exposure of autoradiograms reveals the bands for the second protein (magenta arrow), and supershifts with HA antibody. **e.** Oligo 4 contains two motifs is the only case where the bands corresponding to one (1 TF) or two proteins (2 TFs) could be detected. Gel at the left: Pou5f3 and Nanog can bind both motifs. Gel at the right: 1 µl of Pou5f3 was added to all wells, well 3 also contains 1 µl of Nanog, 2X serial dilutions of Nanog (1/2; 1/4, 1/8, etc) were added to the wells 4 -9. See schematics below for our interpretation of the composition of “1 TF” and “2 TF” bands: we assumed, based on the sequence, that Pou5f3 binds with higher affility to the left motif and Nanog to the right motif. *n.s* – nonspecific band. Uncropped gel images are provided at the end of this file.

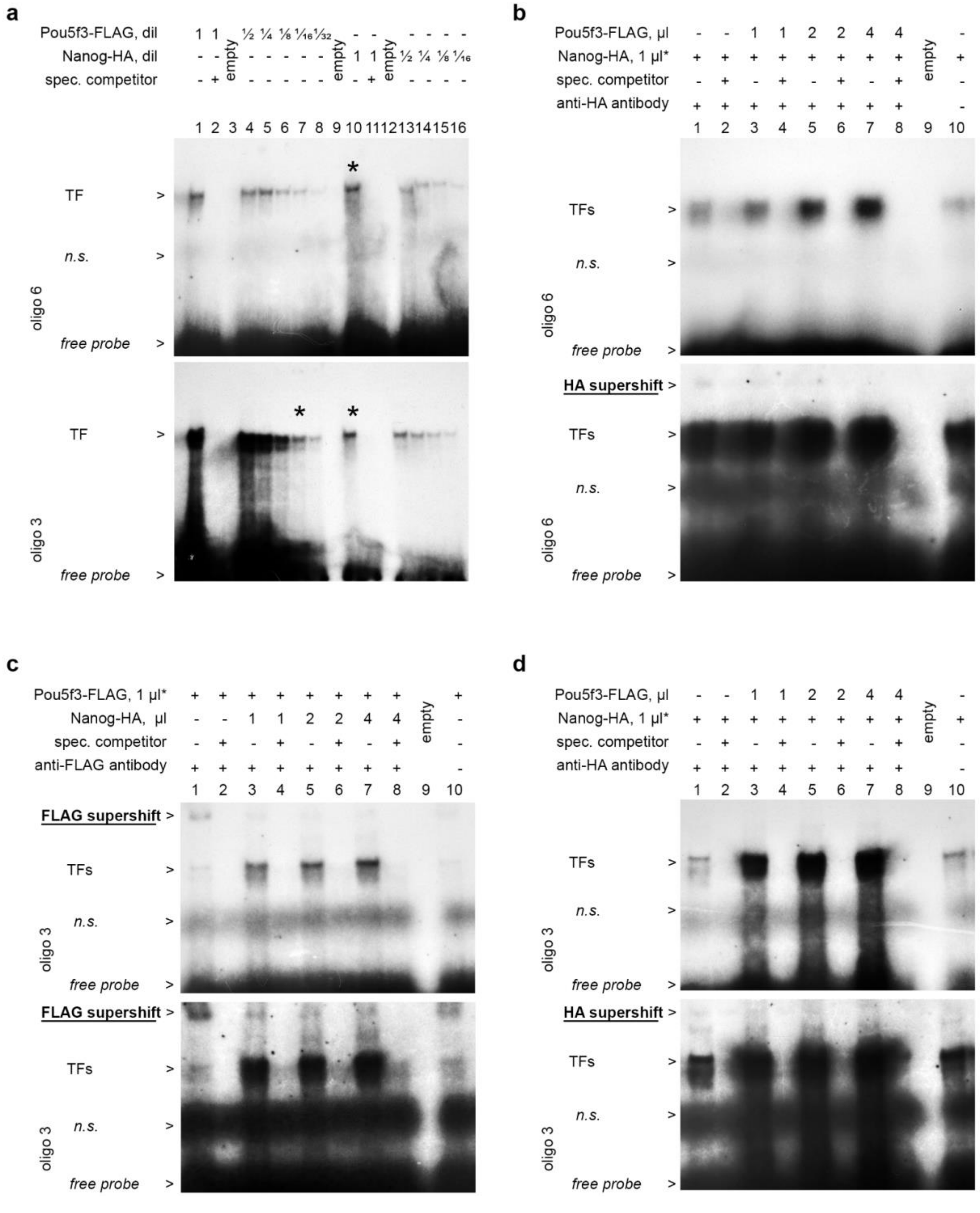

**Figure S7. Pou5f3 and Nanog bind to the common motifs in a mutually exclusive way (related to the main Fig. 5).**

**a**. EMSA. Titration of Pou5f3-FLAG and Nanog-HA protein dilutions against oligo 6 (top) and oligo 3 (bottom). Note that Pou5f3-FLAG and Nanog-HA protein-DNA complexes have similar mobility and cannot be distinguished in the non-denaturing 1xTBE gels. The protein concentrations marked by * were usedfor the first TF in the competition assays shown below. **b-d**.EMSA. Competition assays: the constant amount of first TF was titrated against increasing amounts of the second TF as shown in the legend. Supershifts with anti-FLAG or anti-HA antibody were used to distinguish between Nanog-HA-DNA and Pou5f3-FLAG-DNA complexes. Two images of the same gel are shown: top: 12 h exposure, bottom: 72h exposure. Note that HA supershift is inhibited by increasing concentration of Pou5f3-FLAG (b,d), and FLAG supershift is inhibited by increasing concentration of Nanog-HA (c). *n.s* – nonspecific band. Uncropped gel images are provided at the end of this file.

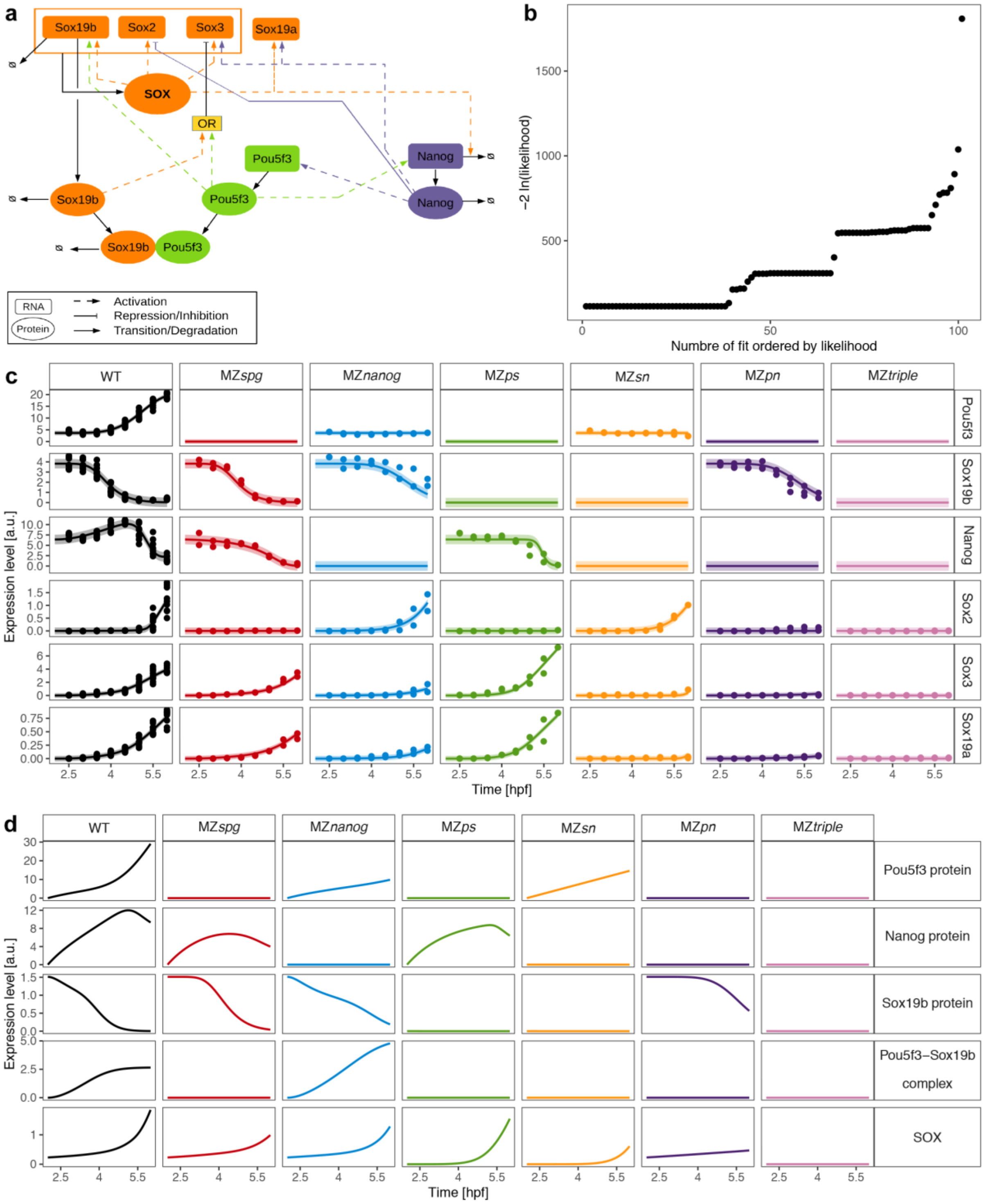

**Figure S8. Core model for dynamics of zygotic genome activators (related to the main Fig. 6).** **a.** Model cartoon of core interactions between SoxB1 family class (orange), Nanog (violet) and Pou5f3 (green). Corresponding mechanistic model equations are summarized in Source Data File 5. **b.** Fit results of the core model are shown as a waterfall plot sorted by their likelihood values. Out of 100 fits, 38 fits ended up at the global optimum. Parameter values of the best fit are given in Source Data File 6. **c-d.** Model trajectories (lines) of RNA targets (c) and protein targets (d) of the best fit are shown together with RNA-seq data (points) used for model calibration. The seven experimental conditions are displayed color coded and in different columns. Different rows correspond to different transcription factors and proteins, respectively. Shaded areas in (c) correspond to uncertainties predicted by the model for each target (compare estimated error parameters in Source Data File 6).

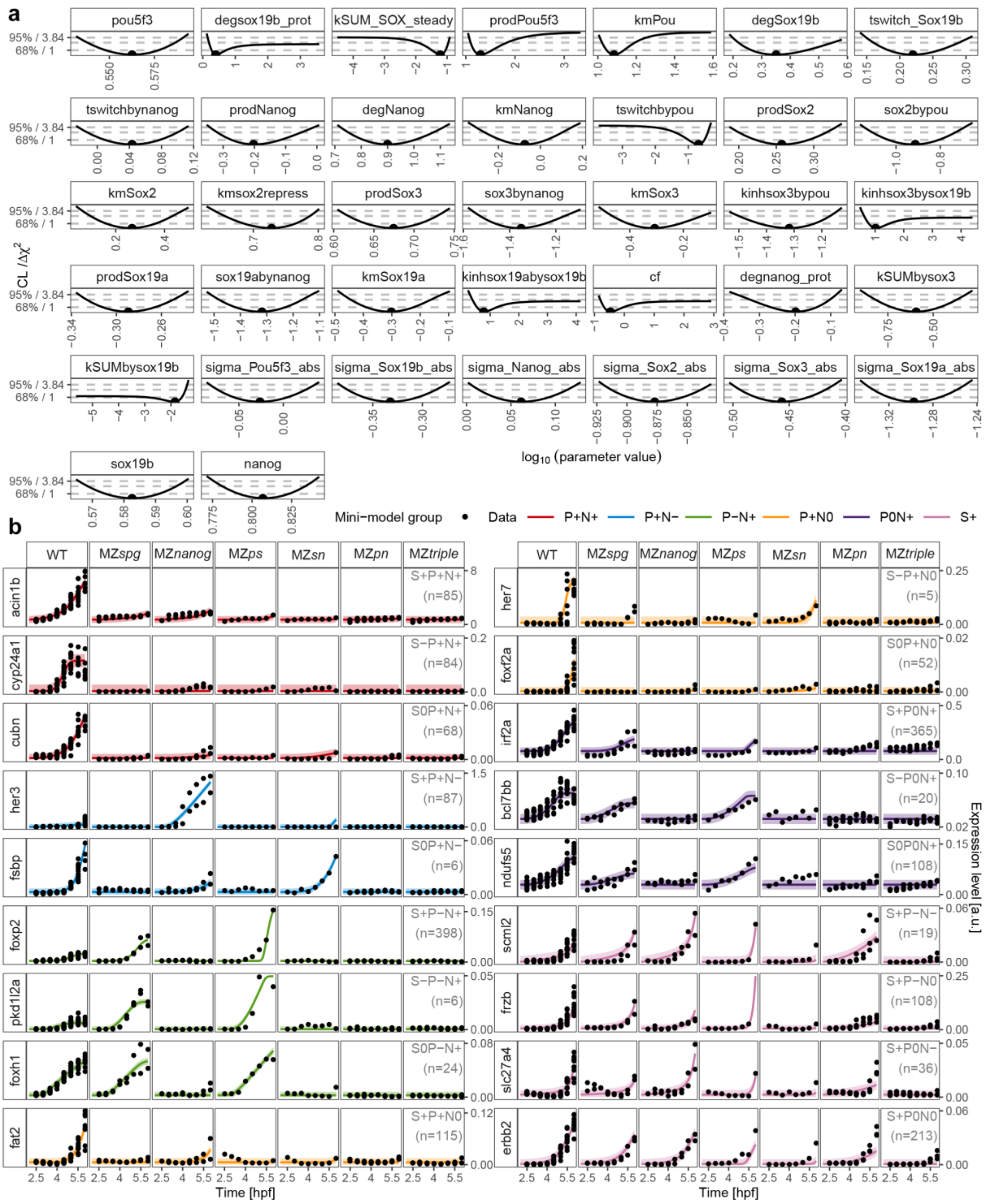

**Figure S9. Identifiability of core model parameters and exemplary mini-model fits (related to the main Fig. 6).**

**a**. Parameter values (dots) of the best fit are shown with their profile likelihoods (lines) for the core model. Likelihood-ratio test based confidence levels for one degree of freedom are shown as dashed grey lines. Parameter values (x-axis) are displayed on log10 scale. Out of 37 estimated parameters, 30 showed parabolic profile likelihoods corresponding to finite confidence intervals, and are thus identifiable[51]. The remaining seven were practically non-identifiable given the available data. The resulting parameter confidence intervals are summarized in Source Data File 6. **b**. For each mini-model one example of the best fit (lines) is shown together with the corresponding RNA-seq data (points). Line colors represent the respective mini-model groups, while n is the total number of transcripts best described by the corresponding mini-model. Columns refer to experimental conditions, and rows indicate different target genes. Model equations representing the different mini-models are summarized in Source data File 7, with bestfit parameter values for the individual target genes shown in Source Data File 7. Mini-model S-P+N- was empty**.**

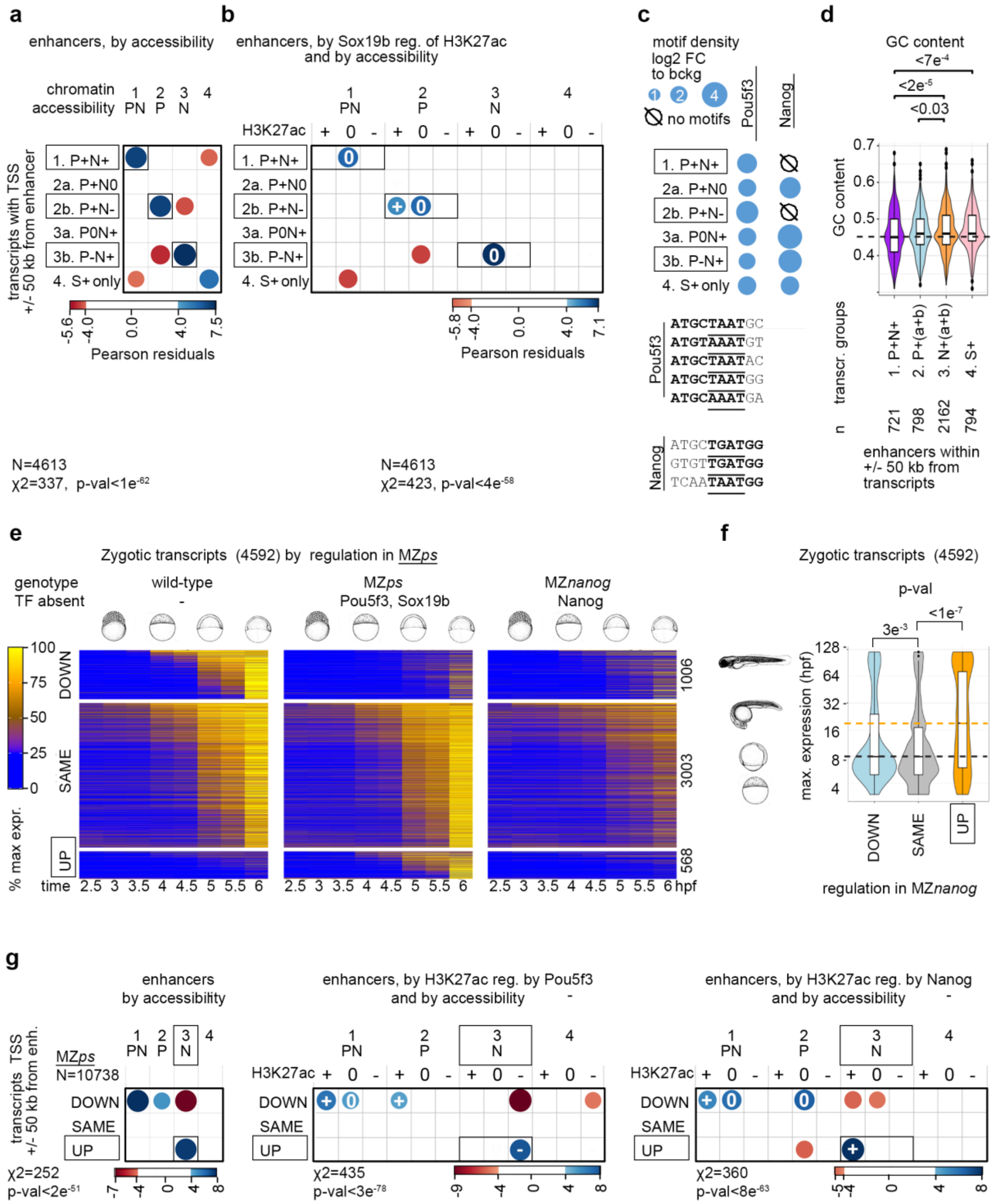

**Figure S10. Pou5f3/Sox19b block transcription activated by Nanog (related to the main Fig. 6 and Fig. 7).**

**a-d.** Validation of transcriptional modelling (“transcriptomics groups”, see main text and main Fig.6). **a, b.** $\chi^2$ tests, positive correlations between the transcriptomics and genomics groups are shown in blue and negative in red. Vertical axis: transcriptomics groups. Direct enhancers were linked to the TSS of zygotic genes within +/-50 kb and sorted according to best fit to one of the six transcriptional model groups. The synergistically regulated targets 1. P+N+ and antagonistically regulated targets 2b. P+N- and 3b. P-N+ are boxed. Horizontal axis: genomic groups. Direct enhancers were sorted to four groups (1.PN, 2.P, 3.N and 4.-) by changes in chromatin accessibility (a), and by H3K27ac regulation by Sox19b (b). +, 0 and – indicate that Sox19b induces H3K27ac, does nothing, or blocks it, respectively. Note that the s+ enhancers are enriched only in the putative regulatory regions of 2b. P+N- transcriptomics group. **c.** Density of Pou5f3- and Nanog- binding motifs from our in-vitro experiments (shown below) differs in the enhancers linked to the different transcriptomics groups. Only exact matches to the sequences shown below were scored. **d**. GC content is significantly lower in the enhancers linked to transcriptomics groups activated by Pou5f3 compared with those activated by Nanog. p-values for significant pairwise differences are shown in Tukey-Kramer test; 1-way ANOVA p-val = $4.8e^{-6}$.

**e**. Zygotic gene expression changes in MZ*ps* double mutant. Heatmap shows normalized expression of three groups of zygotic transcripts ("DOWN", "SAME" and "UP" compared to the wild-type) at eight time points, from pre-ZGA (2.5 hpf) till 6 hpf; numbers of transcripts are shown at the right. **d**. Premature zygotic expression in MZ*ps***.** Expression times in the normal development were compared in "DOWN", "SAME" and "UP", groups; and were significantly higher in "UP" group of transcripts (dotted lines show the median values). P-values in Tukey-Kramer test, 1-way ANOVA p-val < $2e^{-16}$. **e**. $\chi^2$ tests, positive correlations between the transcriptomics and genomics groups are shown in blue and negative in red. Vertical axis: transcriptomics groups. Direct enhancers were linked to the TSS of zygotic genes within +/-50 kb and sorted according to DOWN", "SAME" and "UP" transcriptional groups. Horizontal axis: genomic groups. Direct enhancers were sorted by changes in chromatin accessibility, and by H3K27ac regulation by Pou5f3 or Nanog, as indicated. Note that the Nanog-activated enhancers where H3K27 acetylation is blocked by Pou5f3 (3.N p-n+) are enriched around TSS of genes upregulated in MZ*ps* (boxed). Source data are provided as a Source Data file 2 and as a Source Data file 3. Zebrafish embryo drawings were used with permission of John Wiley & Sons - Books, from "Stages of Embryonic Development of the Zebrafish", Kimmel et al., Developmental Dynamics 203:253-310 (1995); permission conveyed through Copyright Clearance Center, Inc.

Appendix: uncropped gel images for Fig. S6 and S7

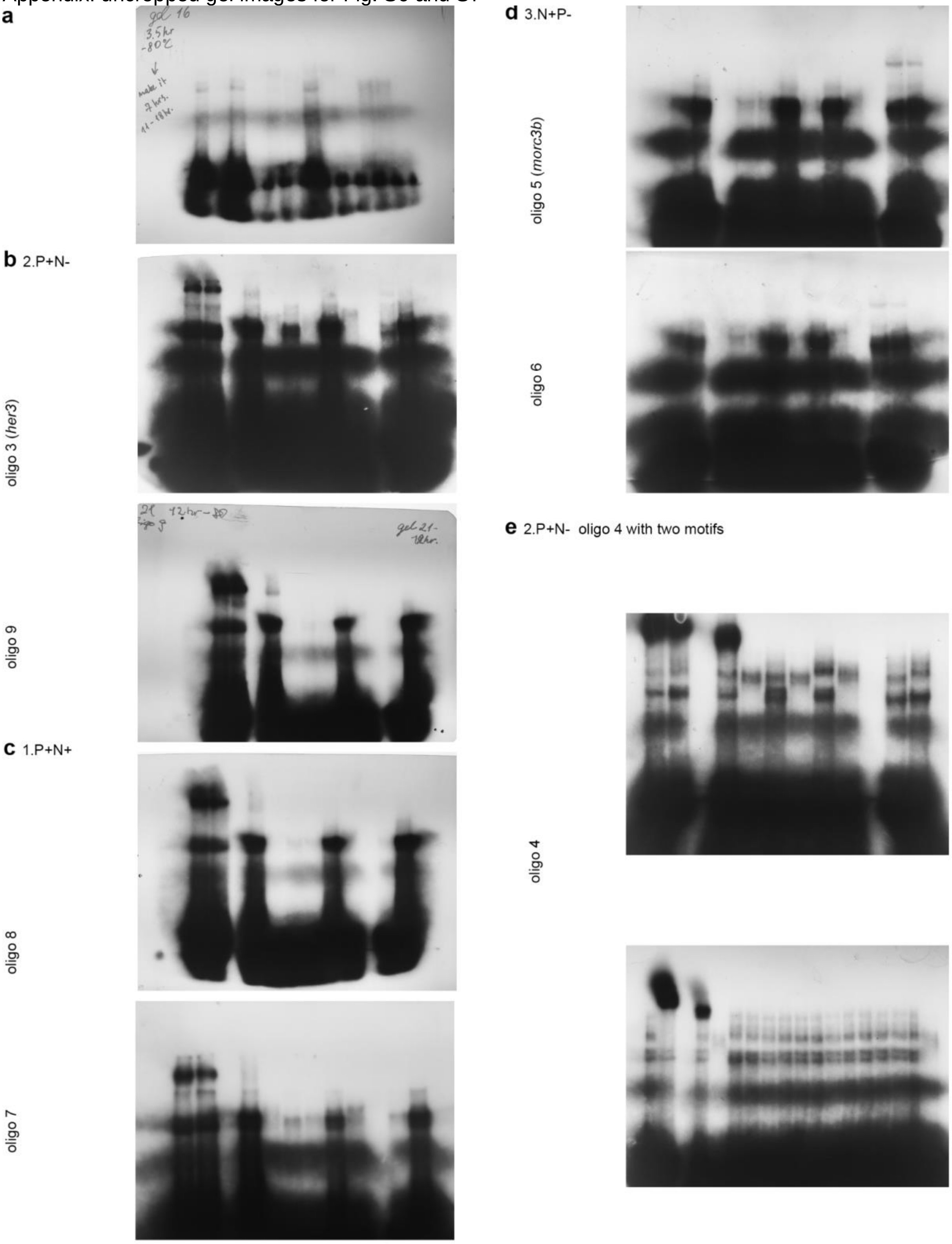

Uncropped gel images for Figure S6.

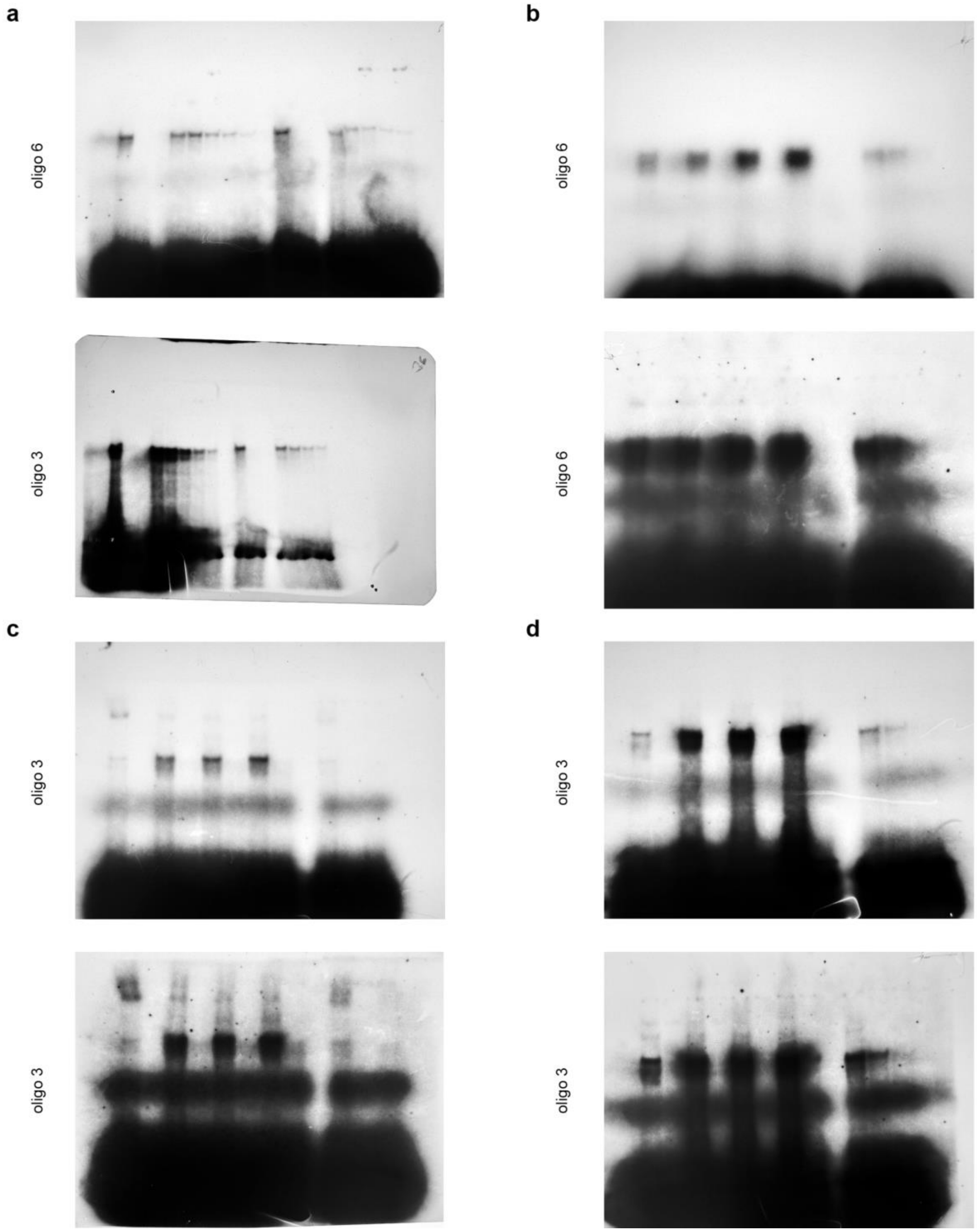

Uncropped gel images for Figure S7

Supplementary Table

| REAGENT or RESOURCE | SOURCE | IDENTIFIER |
|---|---|---|
| Antibodies | | |
| Anti-Histone H3 (acetyl K27) rabbit, 1/100 dilution | Abcam plc., Cambridge, UK | ab 4729 |
| Chemicals, Peptides, and Recombinant Proteins | | |
| cOmplete™, EDTA-free Protease Inhibitor Cocktail | Sigma-Aldrich Chemie GmbH, Germany | 5056489001 |
| SYTOX Green | ThermoFisher SCIENTIFIC | S7020 |
| Critical Commercial Assays | | |
| Agencourt® AMPure® XP Beads | Beckmann Coulter, Krefeld, Germany | A63880 |
| Agilent High Sensitivity DNA Kit | Agilent Technologies, Santa Clara, California, USA | 5067-4626 |
| Agilent RNA 6000 Nano Kit | Agilent Technologies, Santa Clara, California, USA | 5067-1511 |
| Dynabeads® Protein G | invitrogen Dynal AS, Oslo, Norway | 10003D |
| E.Z.N.A® Cycle Pure Kit | Omega Biotek, Norcross, Georgia, USA | D6493-02 |
| Illumina Tagment DNA Enzyme and Buffer Small Kit | Illumina | 20034197 |
| Microcon®-30 Centrifugal Filters | Merck Millipore, Darmstadt, Germany | MRCF0R030 |
| mMESSAGE mMACHINE® SP6 transcription Kit | Ambion | 10086184 |
| NEBNext Ultra DNA Library Prep Kit for Illumina | New England Biolabs, Inc., Frankfurt a.M., Germany | E7370S |
| NEBNext® Multiplex Oligos for Illumina® (Index Primers Set 1) | New England Biolabs, Inc., Frankfurt a.M., Germany | E7335S |
| RNeasy® Mini Kit | QIAGEN, Hilden, Germany | 74104 |
| Quant-iT™ PicoGreen® dsDNA Assay Kit | invitrogen™ Molecular Probes® Waltham, Massachusetts, USA | Q33120 |
| Deposited Data | | |
| Omni ATAC-seq of the wild-type, MZ*sox19b*, MZ*spg*, MZ*ps* zebrafish embryos at 3.7 and 4.3 hpf | Gao et al., 2022 | GEO: GSE188364 |
| Omni ATAC-seq of the wild-type, MZ*nanog*, MZ*pn*, MZ*sn* and MZ*triple* zebrafish embryos at 3.0, 3.7 and 4.3 hpf | this work | GEO: GSE215956 |
| Omni ATAC-seq and H3K27ac ChIP-seq of the wild-type and MZ*nps* zebrafish embryos at 4 hpf | Miao et al., 2022 | SRA: SRP355652 |
| Time-resolved RNA-seq at 8 time points in the wild-type, MZ*sox19b*, MZ*spg*, MZ*nanog*, MZ*ps,* MZ*pn*, MZ*sn* and MZ*triple* zebrafish embryos | this work | GEO: GSE162415 |
| ChIP-seq for H3K27ac in the wild-type, MZ*spg* and MZ*sox19b* zebrafish embryos, 4.3 hpf | Gao et al., 2022 | GEO: GSE143306 |
| ChIP-seq for H3K27ac in the MZ*nanog* zebrafish embryos, 4.3 hpf | this work | GEO: GSE143439 |

| | | |
|---|---|---|
| ChIP-seq for Pou5f3 and Sox2 (SoxB1) TF binding, 5.3 hpf. | Leichsenring et al., 2013 | GEO: GSE39780 |
| ChIP-seq for Nanog TF binding, 4.3 hpf | Xu et al., 2012 | GEO: GSE34683 |
| MNase-seq of MZ*sox19b* embryos, 4.3 hpf | Gao et al., 2022 | GEO: GSE125945 |
| MNase-seq of WT and MZ*spg* embryos, 4.3 hpf | Veil et al., 2019 | GEO: GSE109410 |
| Zebrafish reference genome assembly danrer11/ GRCz11 | Genome reference Consortium | https://www.ncbi.nlm.nih.gov/grc/zebrafish |
| Experimental Models: Organisms/Strains | | |
| Wild-type zebrafish strain AB/TL | ZIRC | ZL1/ZL86 |
| MZ*spg* zebrafish | Lunde et al., 2004 | m793 |
| MZ*nanog* zebrafish | Veil et al., 2018 | m1435 |
| MZ*sox19b* zebrafish | Gao et al., 2022 | m1434 |
| MZ*ps* zebrafish | Gao et al., 2022 | m1434/m793 |
| MZ*pn* zebrafish | this work | m1435/m793 |
| MZ*sn* zebrafish | this work | m1434/m1435 |
| MZ*triple* zebrafish | this work | m1434/m1435/m793 |
| Oligonucleotides | | |
| PCR primer for genotyping Sox19b-f1 5'-ATTTGGGGTGCTTTCTTCAGC-3' | Gao et al., 2022 | no |
| PCR primer for genotyping Sox19b-r1 5'-GTTCTCCTGGGCCATCTTCC-3' | Gao et al., 2022 | no |
| PCR primer for genotyping Nanog1-f1 5'- CAACTTGCCGTCGCTCTAAAC-3' | this work | no |
| PCR primer for genotyping Nanog1-r1 5'- GCTCAGTCTTGTTGTAGGACAG -3' | this work | no |
| PCR primer for genotyping spg-f1 5'-` GTCGTCTGACTGAACATTTTGC -3' | this work | no |
| PCR primer for genotyping spg-r1 5'-` GCAGTGATTCTGAGGAAGAGGT -3' | this work | no |
| PCR primer for H3K27ac ChIP-seq control, positive reference tiparp_f_1 5' CGCTCCCAACTCCATGTATC-3' | Gao et al., 2022 | no |
| PCR primer for H3K27ac ChIP-seq control, positive reference tiparp_r_1 5'-AACGCAAGCCAAACGATCTC-3' | Gao et al., 2022 | no |
| PCR primer for H3K27ac ChIP-seq control, negative reference igsf2_f_2 5'-GAACTGCATTAGAGACCCAC-3' | Gao et al., 2022 | no |
| PCR primer for H3K27ac ChIP-seq control, negative reference igsf2_r_2 5'-CAATCAACTGGGAAAGCATGA-3' | Gao et al., 2022 | no |
| Oligos for the EMSA (5'to 3') | | |
| 1f_EMSA GGGATGGCTTATGCTAATGCCTGATA | this work | no |
| 1r_EMSA GGGTATCAGGCATTAGCATAAGCCAT | this work | no |
| 2f_EMSA GGGTGTTGTAGTGCTAATGTAATGAA | this work | no |
| 2r_EMSA GGGTTCATTACATTAGCACTACAACA | this work | no |
| 3f_EMSA_her3 GGGCATTGACATGCTAATGGAGCTTT | this work | no |
| 3r_EMSA_her3 GGGAAAGCTCCATTAGCATGTCAATG | this work | no |

| | | |
|---|---|---|
| 4f_EMSA_foxb1 GGGTGTCCATGTAAATAGGTGTAAATGAAA | this work | no |
| 4r_EMSA_foxb1 GGGGTTTCATTTACACCTATTTACATGGACA | this work | no |
| 5f_EMSA_morc3b1 GGGGCTGGAGGTGTTGATGGCTGTGT | this work | no |
| 5r_EMSA_morc3b1 GGGACACAGCCATCAACACCTCCAG | this work | no |
| 6f_EMSA_pcdh18b GGGCTGTTCACATGCTGATGGTTCTT | this work | no |
| 6r_EMSA_pcdh18b GGGAAGAACCATCAGCATGTGAACAG | this work | no |
| 7f_EMSA GGGCGCCTGTATGCTAATACAGTCTT | this work | no |
| 7r_EMSA GGGAAGACTGTATTAGCATACAGGCG | this work | no |
| 8f_EMSA GGGCTATGATATGTAAATGTAACAGA | this work | no |
| 8r_EMSA GGGTCTGTTACATTTACATATCATAG | this work | no |
| 9f_EMSA GGGCTTTTTAAATGCAAATGACCTCT | this work | no |
| 9r_EMSA GGGAGAGGTCATTTGCATTTAAAAAG | this work | no |
| 10f_EMSA GGGGTACTGTAATGCAAATGCTGTTG | this work | no |
| 10r_EMSA GGGCAACAGCATTTGCATTACAGTA | this work | no |
| 11f_EMSA GGGGTCTGAAGTGCTAATGAACTGAG | this work | no |
| 11r_EMSA GGGCTCAGTTCATTAGCACTTCAGA | this work | no |
| 12f_EMSA GGGGCTGGAGGTGTTGATGGCTGTTT | this work | no |
| 12r_EMSA GGGAAACAGCCATCAACACCTCCAG | this work | no |
| 13f_EMSA GGGTTTAAATATGAAAGAAATGTGGT | this work | no |
| 13r_EMSA GGGACCACATTTCTTTCATATTTAAA | this work | no |
| 14f_EMSA GGGAATTCATATGCAAATATGTGGG | this work | no |
| 14r_EMSA GGGCCCACATATTTGCATATGAATT | this work | no |
| 15f_EMSA GGGACAGTGCTCAATAATGGCCT | this work | no |
| 15r_EMSA GGGAGGCCATTATTGAGCACTGT | this work | no |
| Software and Algorithms | | |
| Bed Tools | Quinlan and Hall, 2010 | BED Tools in usegalaxy.eu |
| Bowtie2 | Langmead and Salzberg, 2012 | Bowtie2 in usegalaxy.eu |
| DeepTools2 | Ramirez et al., 2016 | deepTools in usegalaxy.eu |

| | | |
|---|---|---|
| DESeq2 | Love et al., 2014 | DESeq2 in usegalaxy.eu |
| FeatureCounts | Liao et al., 2014 | featureCounts in usegalaxy.eu |
| Galaxy server | Afgan et al., 2018 | https://usegalaxy.eu/ |
| geecee utility to calculate fractional GC content of nucleic acid sequences | emboss (version 5.0.0) | geecee in usegalaxy.eu |
| GREAT: Genomic Regions Enrichment of Annotations Tool, version 3.0.0 | Hiller et al., 2013 | http://great.stanford.edu/great/public-3.0.0/html/ |
| *In-vitro* nucleosome prediction program | Kaplan et al., 2009 and https://github.com/bgruening/galaxytools | Nucleosome Predictions in usegalaxy.eu |
| MACS2 | Ferg et al., 2007 | MACS2 callpeak and MACS2 bdgpeakcall in usegalaxy.eu |
| R: Version 3.6.1 | R Core Team (2019). R: A language and environment for statistical computing. R Foundation for Statistical Computing, Vienna, Austria. | https://www.R-project.org/. |
| RNA Star | Dobin et al., 2013 | RNA Star in usegalaxy.eu |
| RNA-sense | Gao et al., 2022 | https://bioconductor.org/packages/release/bioc/html/RNAsense.html |
